# Revisiting co-expression-based automated function prediction in yeast with neural networks and updated Gene Ontology annotations

**DOI:** 10.1101/2025.03.27.645865

**Authors:** Cole E. McGuire, Matthew A. Hibbs

## Abstract

Automated function prediction (AFP) is the process of predicting the function of genes or proteins with machine learning models trained on high-throughput biological data. Deep learning with neural networks has become the dominant machine learning architecture of contemporary AFP models. However, it is unclear what difference exists between neural networks and previous machine learning architectures for AFP. Therefore, we created a model of AFP in yeast using neural networks that is trained on gene co-expression data to predict Gene Ontology (GO) labels. When trained on the same input data, we found that our model outperforms two other experimentally-validated co-expression-based AFP models using other machine learning techniques (Bayesian networks and adaptive query-driven search) when predicting individual genes involved in mitochondrion organization. In particular, we found our neural network model better distinguished mis-annotated negatives in its training data. Finally, we quantified how differences in the gene expression data and Gene Ontology annotations affect the performance of our model across each of its predicted GO terms. Our results suggest that neural networks are more performant and robust to GO mis-annotations compared to other machine learning architectures for co-expression-based AFP of some biological processes.

## 1. Introduction

Automated function prediction (AFP) is the process of predicting the function of genes or proteins by learning patterns in high-throughput biological data using machine learning. Ideally, AFP methods make predictions that can inform experimental biology to increase the rate that researchers can make novel biological discoveries. The history of AFP shows a trend of using increasingly sophisticated machine learning models and making predictions in increasingly complex organisms. One conception of AFP uses machine learning models trained on high-throughput biological data to predict gene or protein function in terms of Gene Ontology labels [1]. The first iterations of this approach occurred in 2003 and 2004 using models built with Bayesian networks. These models combined high-throughput gene co-expression, protein interaction, and genetic interaction data along with expert-curated priors to predict gene function in *Saccharomyces cerevisiae* [2,3].

SPELL, MEFIT, and bioPIXIE were subsequent models of automated function prediction in *S. cerevisiae* that leveraged gene co-expression data and innovated on many of the ideas found in prior work [4–6]. MEFIT and bioPIXIE employed naive Bayesian networks while SPELL used a directed query search through gene expression datasets. These were among the first AFP methods to use some conception of model training, since MEFIT and bioPIXIE replaced expert-curated model priors with priors learned from training data, and SPELL dynamically generated model weights at runtime in response to each query. These three methods were trained on similar input data and were experimentally validated in a systematic study of mitochondrion organization and biogenesis in yeast. This led to the discovery of over 100 genes involved in that process [7–9], and demonstrated that AFP can be useful for directing experimental biology. At the same time, models were developed that used more sophisticated machine learning methodologies, such as support vector machines [10–12]. Additionally, methodologies began to branch out to predicting gene and protein function in organisms other than yeast [13,14], and predictions were extended beyond GO annotations to predict other genomic features, such as knockout phenotypes and gene-disease associations [15–17].

In the past decade, much of the cutting edge research in machine learning, and its applications to biology, have transitioned towards the use of neural networks. Neural networks transform input data through layers of “artificial neurons” to produce an output prediction. Neural networks quantify the discrepancy (or loss) between the predicted output and the target output of labeled training examples using a loss function. Using this loss value, neural networks can automatically update their own parameters to make more accurate predictions via a process known as “backpropagation.” Deep neural networks (those with several layers of artificial neurons) are responsible for breakthroughs in numerous problem domains including image recognition and natural language processing due to their ability to learn complex hierarchical, non-linear representations of data. However, the complexity of the functions learned by deep neural networks makes it challenging for humans to understand the rationale behind how these model’s make their predictions. Improving the explainability of neural network outputs and finding ways to interpret the inner workings of these models is an ongoing area of research [18].

Advancements in deep learning with neural networks are responsible for a paradigmatic shift in how researchers approach automated function prediction. Deep neural networks have displayed a marked increase in predictive accuracy over past AFP models [19]. Several variants of neural networks have shown to be particularly well suited for modeling complex relationships in various modalities of biological input data; some notable variants include convolutional neural networks, graph neural networks, and transformers [19]. Furthermore, neural networks have been associated with a trend in AFP towards predicting function from primarily a protein’s amino acid sequence (in the absence of species information) [20–22].

The rise of deep neural networks for AFP has exacerbated a trend where AFP models lose the attention of researchers once they have been surpassed in predictive accuracy by newer, state-of-the-art methods. This trend of chasing predictive accuracy above all else has generated a gap in knowledge; we lack an understanding of how the predictions of neural network AFP models differ from previous methods beyond having a higher predictive accuracy on benchmarks. Additionally, most AFP models lack systematic experimental validation beyond evaluating performance on newly annotated proteins (that were not present when the model was trained) [1]. Therefore, it is unclear how most AFP models differ when it comes to directing novel biological experimentation.

In this paper, we attempt to bridge this knowledge gap by directly comparing neural networks to previous, experimentally-validated machine learning techniques at the task of predicting GO biological process labels from high-throughput gene co-expression input data in yeast. We built a model of AFP using linear neural networks that predicts 79 GO biological process labels from co-expression data of pairs of genes. We trained our model with the same gene expression data and GO labels that MEFIT and SPELL were trained with in 2007. When comparing each model’s ability to predict genes involved in mitochondrion organization, we found our model is more performant than MEFIT and SPELL, in part due to its ability to better distinguish mis-annotated negatives in its training data. Finally, we trained our model with current-day gene expression data and GO labels to assess how the accumulation of more input data and higher quality training labels affects the performance of our model.

## 2. Methods

Our model trains fully-connected linear neural networks to predict the functional relationship between pairs of genes in the context of 79 Gene Ontology biological processes (Fig 1). As input, our model uses the correlations between the gene expression profiles of pairs of genes in 113 microarray assay datasets. Once trained, our model uses its predictions to construct a functional relationship graph for each GO term that can be converted into experimentally-testable single gene functional predictions. Our initial model is trained using the same Gene Ontology annotations and gene expression datasets that SPELL and MEFIT trained with as of April 15th, 2007.

**Fig 1.**
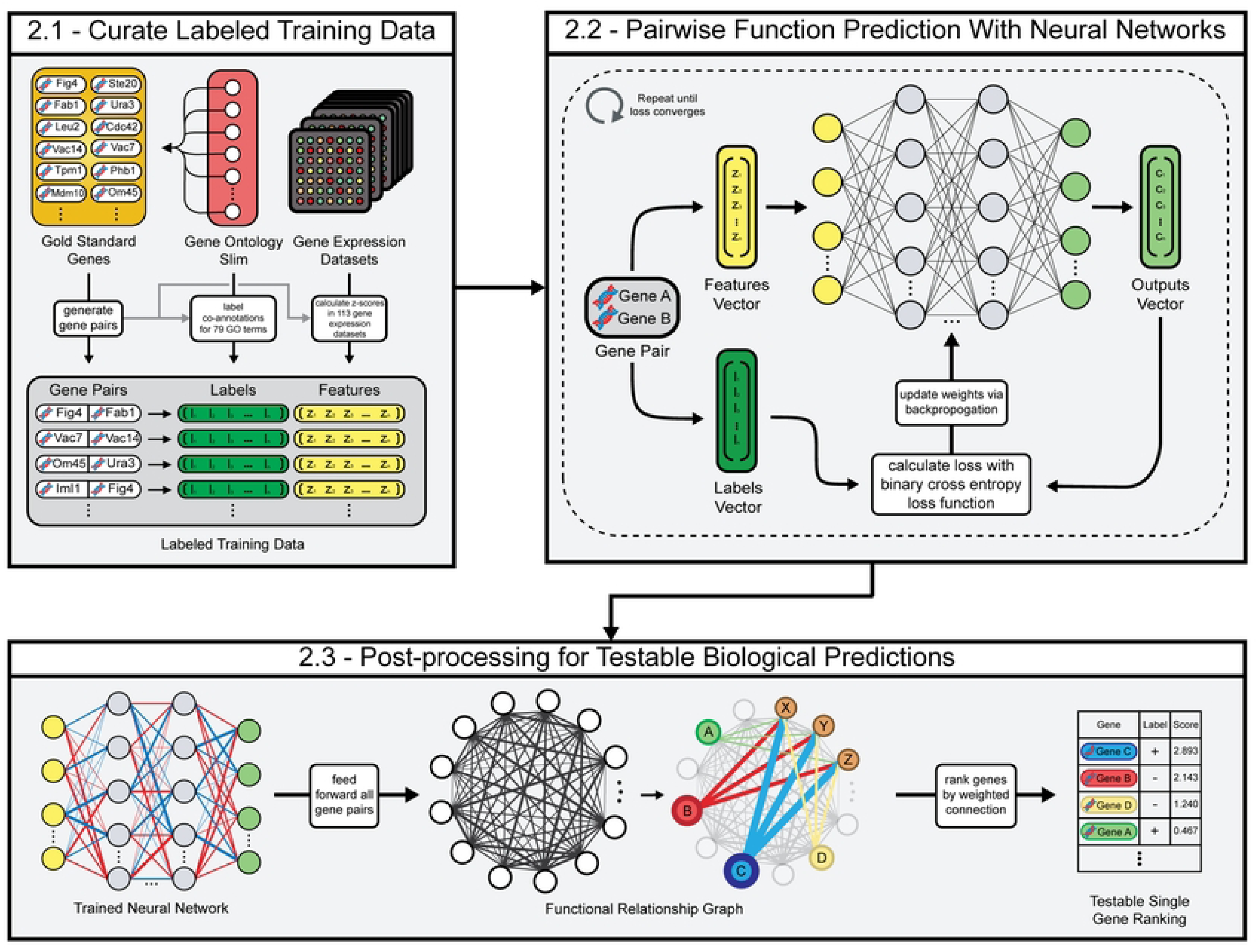
**Overview of methodology**. (A) Training data is created by calculating pairwise gene expression similarities in 113 gene expression datasets and by curating a set of “gold standard” training labels for genes with known functions. (B) Linear neural networks are trained to predict the Gene Ontology co-annotations of pairs of genes given feature vectors of their gene expression similarity. (C) Our method translates pairwise functional relationship predictions into testable single gene rankings for 79 biological processes.

### 2.1 Curation of labeled training data

The input data for our model represents each pair of genes as a vector of 113 z-scores derived from Pearson correlations across experimental conditions in each of 113 gene expression datasets. To train our model, we curated training labels for gene pairs with known functional relationships and gene pairs thought to be unrelated. Positive and negative labels are determined for each of the 79 predicted GO terms based on each gene pair’s co-annotation to a term, as detailed below.

#### 2.1.1 Gene expression compendium curation

In order to accurately compare between approaches, our model uses the same gene expression compendium as MEFIT and SPELL, which we obtained from the *Saccharomyces* Genome Database (SGD) [23,24]. This compendium consists of 113 gene expression datasets publicly available and curated by SGD as of April 15th, 2007 and comprises 1842 experimental conditions. These raw gene expression levels are processed for use as input data by normalizing Fisher transformed Pearson correlations between pairs of genes for each of the 113 datasets in the compendium. For each gene expression dataset, the Pearson correlation between a pair of genes’ expression values is calculated as:

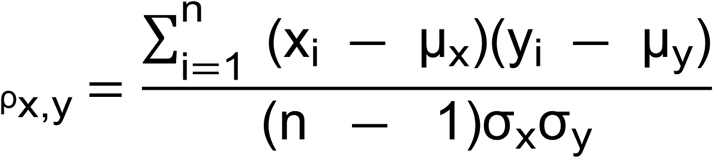

where *x* and *y* are the gene expression vectors of size *n* from a given dataset, *μ_x_ and μ_y_* are their means, and *σ_x_* and *σ_x_* are their standard deviations. In the case of missing measurements, gene expression vectors *x* and *y* only contain the experimental conditions that have observations for both gene x and gene y (Further details and other minor adjustments are described in Supplementary Methods). As the distribution of Pearson correlations can vary greatly between gene expression datasets, the Fisher transformation is applied to improve comparability across datasets [25]:

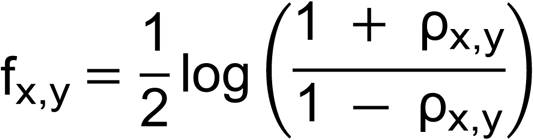

After Pearson correlations have been Fisher transformed, the resulting values are normalized by the following equation to produce a standard normal score (z-score) for each pair of genes:

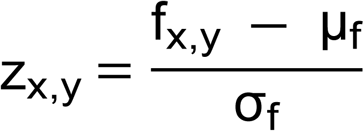

where *μ_f_* and *σ_f_* are estimated mean and standard deviations calculated using a subset of 200,000 random Fisher transformed Pearson correlations from that dataset. The resulting distribution is approximately standard normal [∼N(0,1)]. In all, each value in each dataset is a z-score that represents the number of standard deviations away a gene pair’s gene expression correlation is from the dataset’s mean of pairwise correlations [5]. This is the same preprocessing and normalization approach MEFIT and SPELL used for their gene expression compendiums [4,5].

#### 2.1.2 Gold standard curation and cross validation

To train our model, we need training labels consisting of gene pairs known to be functionally related (positive examples) and gene pairs thought to lack functional relationships (negative examples). To derive these labels, we first curate a set of single genes that have known functions, which we will call our “gold standard” genes. We defined genes with known functions as genes annotated to at least one of a set of 79 biological process Gene Ontology terms curated by experts as being informative enough to direct laboratory experiments [26]. These GO terms form a subset of biological process terms (distributed as a GO slim), and we utilized annotations to this subset from January 1st, 2007 in order to enable comparison to prior methods trained at that time. We further filtered the GO slim to only include terms with a sufficient number of annotations for model training (n >= 10). Genes not annotated to any of these 79 GO terms were considered functionally uncharacterized.

Our model uses 4-fold cross validation to assess overfitting and make its final predictions. The gold standard genes are randomly divided into four independent subsets. Each “fold” of the model will use three of these subsets as training genes while holding out the remaining subset as testing genes. Each fold generates a set of related (positive) gene pairs and unrelated (negative) gene pairs from the training genes or testing genes. Related pairs are those that are co-annotated to at least one of the 79 GO terms. Unrelated pairs are those whose smallest co-annotated GO term contains at least 10% of the genes in the yeast genome, indicating that each gene has a known function, and those functions are different from each other. This approach is not a perfect measure, since it assumes that two genes with known, different functions are unlikely to share a functional relationship. Note that to avoid data leakage and circularity, gene pairs used as training examples are only pairs where both genes are in the training gene set, and testing examples are only pairs where both genes are in the testing geneset [27,28].

#### 2.1.3 Features and training labels

Our model represents each gene pair as a vector of input features associated with a vector of training labels. The input features for each gene pair is a vector of 113 z-scores derived from the gene expression datasets, and the training labels are one-hot encoded vectors representing the co-annotation of a gene pair to 79 Gene Ontology terms. For each GO term, a gene pair is labeled positive (encoded as 1) if the genes are co-annotated to that GO term and negative (encoded as 0) otherwise.

### 2.2 Pairwise function prediction with neural networks

Our model trains four cross-validated, fully-connected linear neural networks to predict 79 GO term labels from feature vectors of gene expression z-scores. The performance of each fold’s neural network is measured for each GO term, and folds are compared to each other to evaluate the consistency of the cross-validated neural networks. After the model’s performance is evaluated, the model feeds all gene pairs through the neural networks to produce a fully-connected functional relationship graph for each GO term, where nodes correspond to genes, and edge weights correspond to the neural network output confidence predictions.

#### 2.2.1 Neural network architecture and training loop

These data features and labels are used to train fully-connected linear neural networks to predict a gene pair’s co-involvement in each of the previously described 79 GO terms. The neural network was implemented with PyTorch (PyTorch 1.10.2; Python 3.9.12). The network consists of an input layer with 113 neurons and an output layer of 79 neurons. The number and size of the model’s neural networks’ hidden layers is flexible. The model that we use to make our final comparisons against SPELL and MEFIT has three hidden layers with 500, 200, and 100 neurons respectively. Each hidden layer applies a ReLU activation function. At the beginning of the training process, weights are initialized by sampling a uniform distribution U 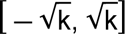 where *k* = 1/ output size of a given layer.

Our model represents the problem of gene pair function prediction as a multilabel prediction problem by using a binary cross entropy loss function; the prediction of each GO term is treated as an independent binary classification task. Our model is trained with stochastic gradient descent; during each iteration of the training loop, a mini-batch of 50 gene pairs is sampled from the training data. This mini-batch consists of 25 gene pairs sampled from the related pairs and 25 gene pairs sampled from the unrelated pairs. After the mini-batch is fed forward through the model, the binary cross entropy loss between the network’s output and training labels is used to update the neural network’s weights via back propagation. The training loop runs for enough iterations for the loss of each mini-batch to converge, which we empirically found to be at least 600,000 iterations. Each network is trained with a learning rate of 0.01 and momentum of 0.9 as hyperparameters.

#### 2.2.2 Performance evaluation

The performance of each cross-validated neural network is measured with receiver operating characteristic (ROC) curves, which show the tradeoffs between the true positive rate (TPR) and false positive rate (FPR). Once a network has been trained, the performance of each of the 79 output GO terms is measured by ranking the prediction of positive pairs and negative pairs derived from the fold’s testing genes or training genes. For a given GO term, positive pairs are gene pairs that are co-annotated to the GO term and negative pairs are pairs of genes where neither gene is annotated to the GO term. A random subset of up to 10000 positive pairs and a random subset of 10 times the number of the positive pairs are evaluated. After these testing pairs have been fed forward through the network, each pair is ranked by its output value for the GO term being evaluated, and a ROC curve is generated using this ranking.

The performance for each GO term is summarized with the average area under the ROC curve (AUC) across all four folds of a model. The overfitting of a single GO term is assessed by comparing the average testing and training data AUCs of that term across each fold of the model.

#### 2.2.3 Generation of functional relationship graphs

Once the networks have been trained, functional relationships are predicted for all possible gene pairs in the yeast genome. The final scores for each gene pair are calculated as the average outputs of the neural network folds that held out either of the genes in a pair. This is done so that only independent network folds contribute to the final score of a pair, meaning that folds are only used to predict pairs they were not trained on. Since pairs that contain uncharacterized genes (which are not included in the gold standard) are held out of all folds’ training data, these pairs’ final scores are calculated as the average predictions of all folds of the model.

Once all pairwise scores between genes have been calculated, a functional relationship graph is constructed for each GO term using each gene pair’s score for that GO term.

A functional relationship graph for a GO term is a weighted graph where the vertices represent genes and the weighted edges represent the predicted confidence of a pair of genes sharing a functional relationship for that GO term.

### 2.3 Post-processing for testable biological predictions

The aggregated outputs of our neural network model are functional relationship graphs for each predicted GO term that contain information about how confident our model is that two genes are functionally related in the context of each respective GO term. Importantly, our predicted GO terms were collected from a list of biological processes experts deemed to be informative for directing experimental biology [26]. While our model’s pairwise predictions contain a wealth of potentially novel biological information, we expect that they are also fairly noisy. We can reduce this noise by aggregating pairwise predictions into predictions of the functions of single genes. Furthermore, this allows us to compare against the performance of other automated function prediction models (such as MEFIT and SPELL) that also make single gene functional predictions. We apply a process similar to MEFIT and SPELL [7,9], where we process our functional relationship graphs to generate testable predictions by ranking individual genes by their predicted involvement in a GO term as measured by the sum of their connections to other genes annotated to that term (Figure 2).

**Figure 2.**
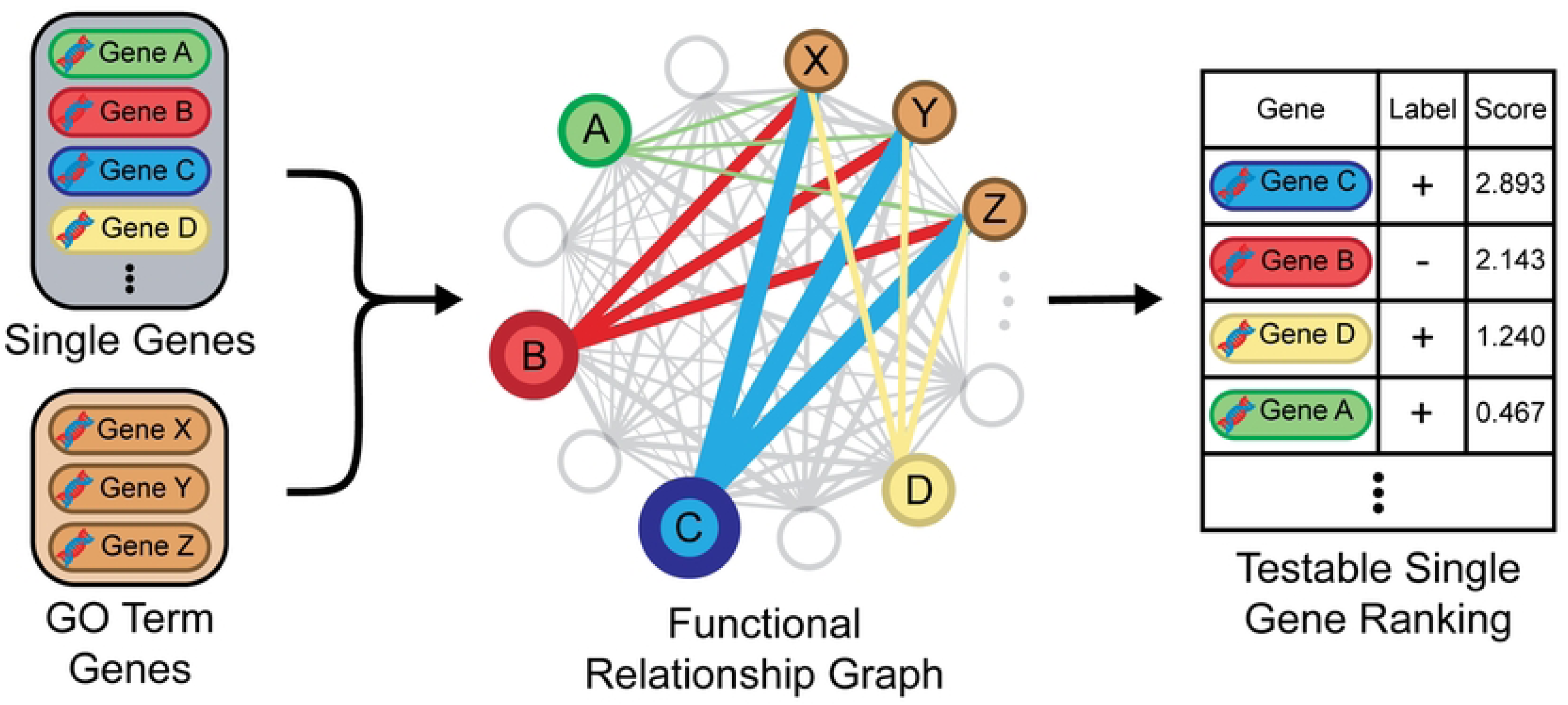
**Diagram illustrating process of generating testable predictions from predicted pairwise relationships**. For each GO term, every gene in the yeast genome is ranked by the weighted average of its connections to genes annotated to that GO term in that GO term’s functional relationship graph.

To predict an individual gene’s involvement in a GO term’s biological process, we use the confidence of the gene’s predicted relationship to other genes annotated to that term. For each GO term’s functional relationship graph, each gene in the yeast genome is ranked by its average weighted connection to all other genes annotated to that GO term. Therefore, for each GO term, our model produces a ranked list of individual genes where the top ranked genes are predicted to be more likely to be involved in that biological process. The performance of the model is measured by generating ROC and precision-recall curves from these single gene ranking [29]. Genes annotated to a GO term are labeled positive, all other genes in the gold standard set are labeled negative, and all uncharacterized genes are unlabeled and do not affect the performance evaluation.

## 3. Results

### 3.1 Model performance

We evaluated the performance and overfitting of our neural network model by comparing the pairwise training AUC, pairwise testing AUC, and single gene AUC for each of the 79 predicted GO terms. We plotted each GO term’s testing AUC against its training AUC with the size and color of each point being proportional to its GO term’s single gene AUC (Fig 3). The overfitting for each GO term (measured as the difference between its training and testing AUC) is represented by its point’s distance from the black diagonal line. We found that a GO term’s pairwise testing AUC was positively correlated with its single gene AUC (r = 0.847; p << 0.001) and negatively correlated with the overfitting (r = -0.859; p << 0.001). Together, this indicates that while our model does suffer from some overfitting, it is significantly less for the GO terms that have high testing AUC performance.

**Fig 3.**
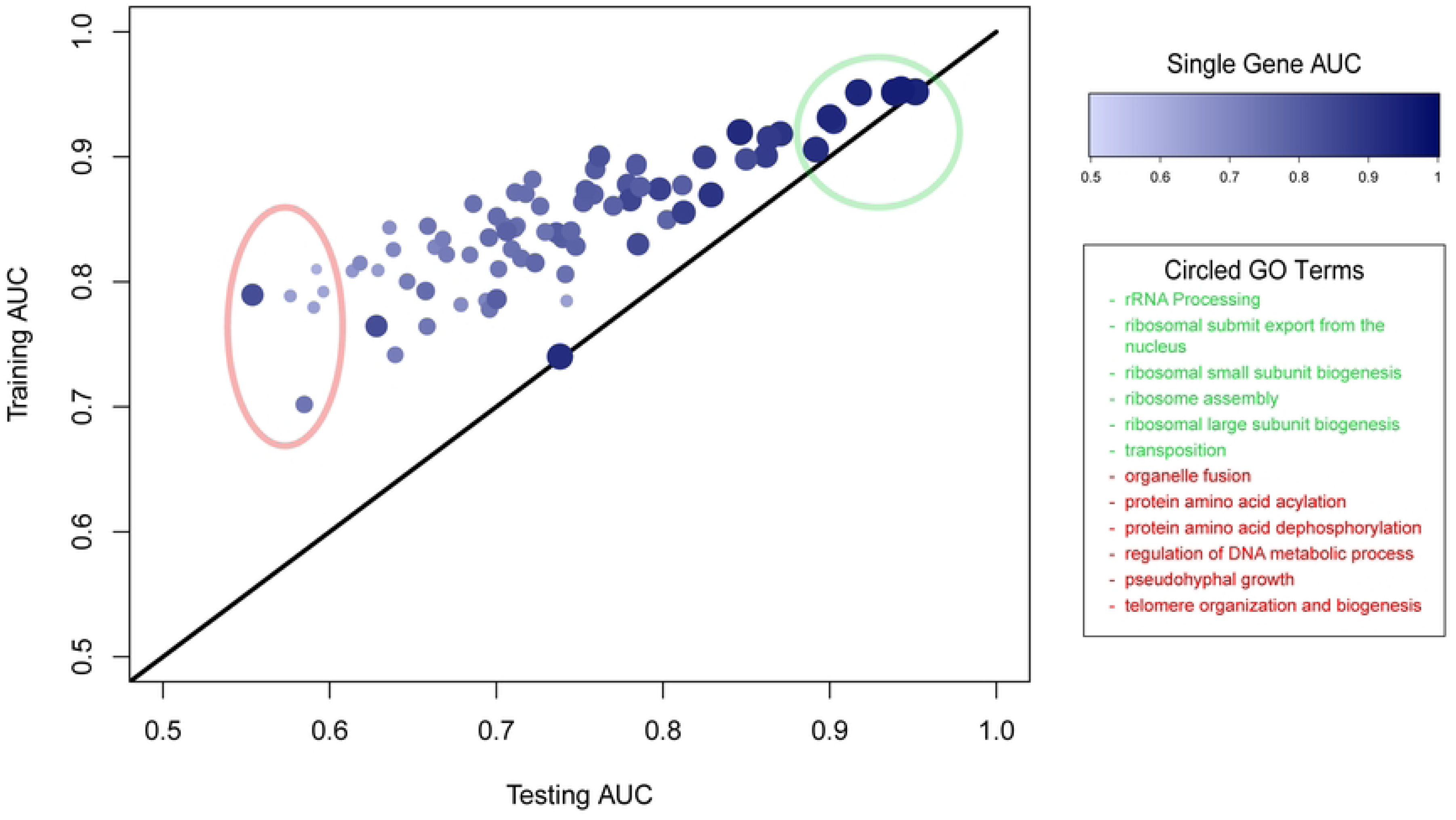
Comparison of performance of neural network model for all 79 output GO terms. For each GO term, the term’s testing AUC is plotted against its training AUC, and the size and color of the point is proportional to the AUC of the GO term’s single gene ranking. The black diagonal line shows where the testing AUC and training AUC are the same, which indicates there is no overfitting. The red circle and text denoted the six GO terms with the worst testing AUC, and the green circles and text denoted the six GO terms with the best testing AUC.

There is great variation in the testing AUC performance and overfitting across all predicted GO terms of the model. Our expectation is that this is the result of our input gene expression data being fundamentally more informative predicting some biological processes compared to others. To examine this claim, we first analyzed the top six best performing and the top six worst performing GO terms by testing AUC, which are the terms circled in green and red respectively (Fig 3). We found that the highest ranking terms were enriched for processes related to ribosome biogenesis, which is a process whose genes are highly co-regulated by gene expression [30]. Conversely, we expected that the weakest performing terms likely lacked strongly coordinated gene co-expression among most pairs of genes annotated to these processes among our data compendium. As such, we found that the lowest ranking terms were pseudohyphal growth (a process not commonly occurring in laboratory yeast strains ([31])) and terms encompassing a diverse set of biological pathways (organelle fusion, protein acylation and dephosphorylation, etc.).

These findings align with our expectation that the testing AUC performance and overfitting for each GO term are correlated with the extent to which a GO term’s genes are co-regulated at the level of gene expression. To evaluate this quantitatively, we examined the difference in co-expression distributions for co-annotated gene pairs to the background distribution for all gene pairs. For each GO term, we derived a probability distribution, *T,* of the average z-scores (across our 113 gene expression datasets) by randomly sampling 200,000 gene pairs co-annotated to that GO term. Additionally, we created a background probability distribution, *B,* by sampling the average z-scores for 200,000 of all the training gene pairs. For each GO term, we measured the Kullback-Liebler (KL) divergence between the term’s probability distribution and the background probability distribution as D_KL_(*T,B*). KL divergence measures the extent of discrepancy between two distributions by quantifying the out on information lost by approximating one distribution with another [32]. The KL divergence of each term was positively correlated with the testing AUC of the term (r = 0.735; p << 0.001) and negatively correlated with the term’s overfitting (r = -0.660; p << 0.001) (S2 Fig). This indicates that as a GO term’s pairwise co-expression differs more from the background co-expression distribution, the GO term tends to make more accurate predictions and suffer from less overfitting. Therefore, these findings support our expectation that our model performance for a given GO term is correlated with the extent to which the term’s genes have coordinated co-expression.

### 3.2 Comparing mitochondrion organization predictions to SPELL and MEFIT

We compared the performance of our neural network model, SPELL, and MEFIT by comparing the ROC curves generated by each model’s rankings of single genes predicted to be involved in mitochondrion organization using Gene Ontology labels from January 1st, 2007 (Fig 4). We found that our neural network model performed better than both SPELL and MEFIT in the TPR-FPR tradeoff as reflected by over a 0.04 increase in AUC compared to both models.

**Fig 4.**
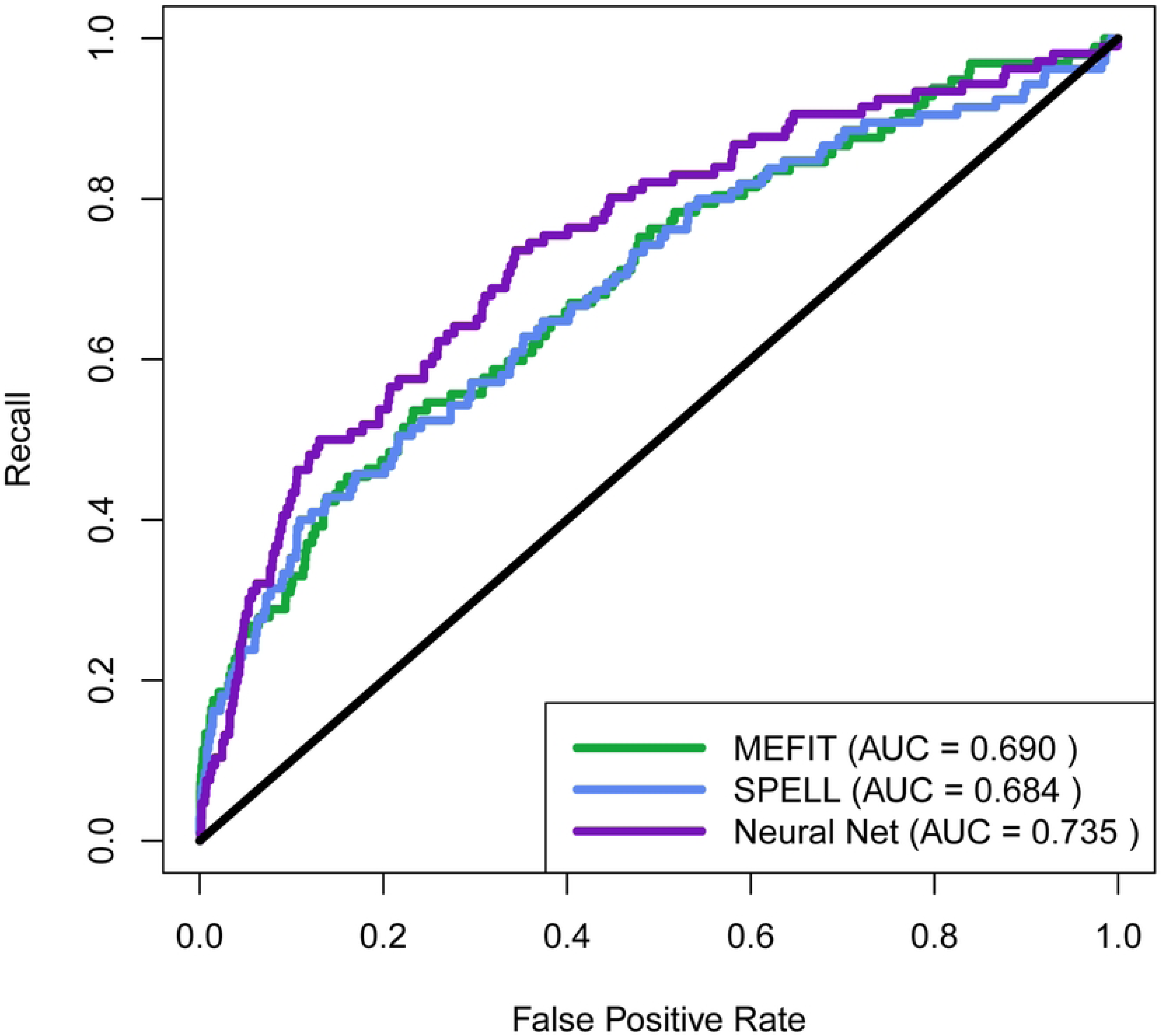
Comparison of ROC Performance to MEFIT and SPELL. ROC curve comparing tradeoff between recall and false positive rate for rankings of genes predicted to be involved in mitochondrion organization for MEFIT, SPELL, and our neural network model. GO labels used in ROC curve calculations were derived from the January 1st, 2007 Gene Ontology.

We also assessed the precision of each model on their top predictions, since those predictions are the most ripe for experimental validation. Using the same rankings mentioned above, we generated precision-recall curves for each model’s rankings (Fig 5.A). We found that the neural network model had worse precision at low recall (0-0.2) compared to SPELL and MEFIT when using labels derived from the 2007 Gene Ontology, despite our model’s overall improvement in ROC performance. In order to assess which model has better precision on top predictions in terms of our current understanding of biology, we relabeled the aforementioned rankings from each model using labels derived from June 16th, 2022 GO annotations. Using these modern labels, we found that our neural network model had higher precision than SPELL and MEFIT at low recall (Fig 5.B).

**Fig 5.**
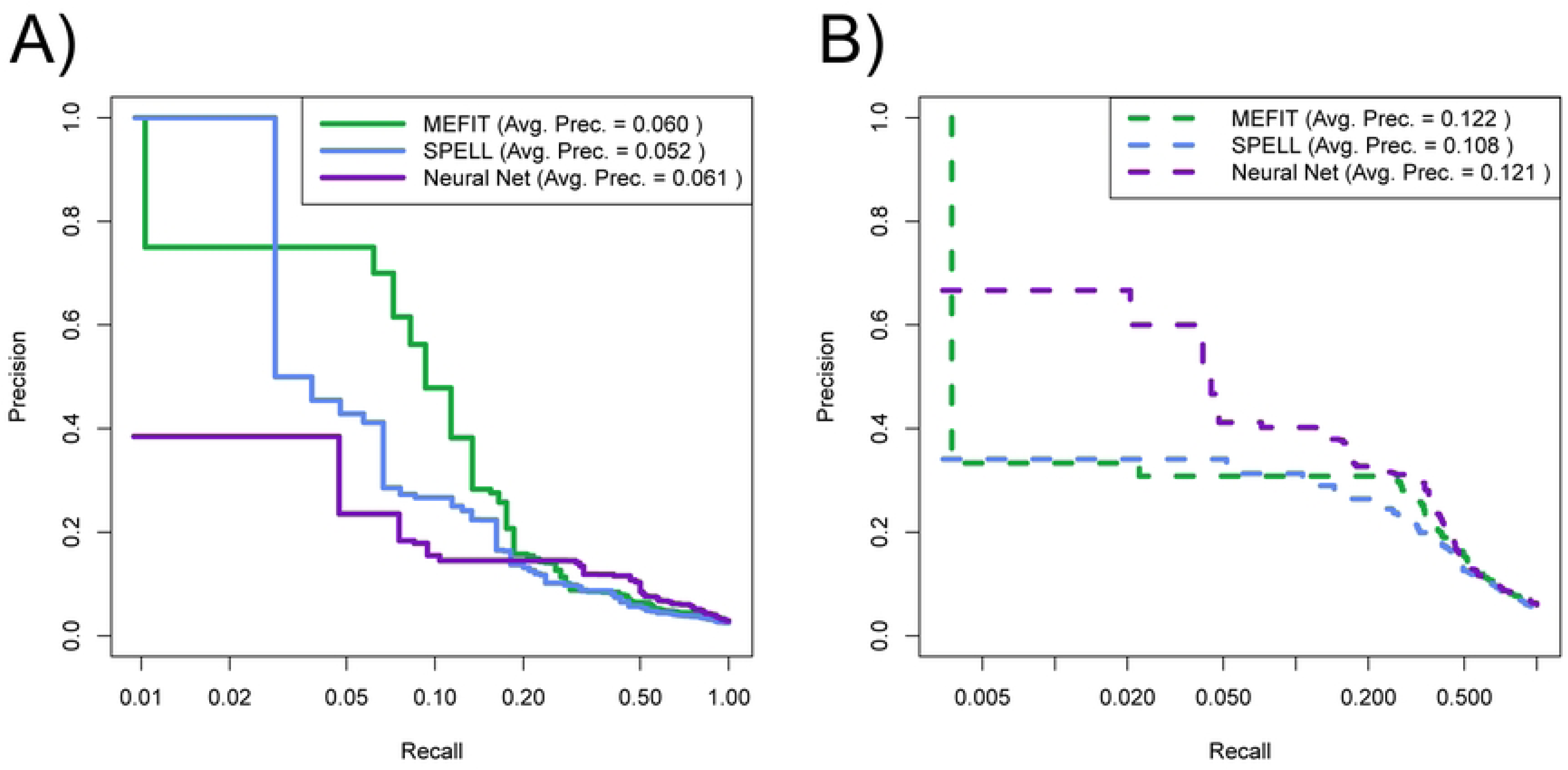
Comparison of precision-recall tradeoff with MEFIT and SPELL. Precision-recall curves with logarithmic x-axes comparing the tradeoff between precision and recall for rankings of genes predicted to be involved in mitochondrion organization for MEFIT, SPELL, and our neural network model trained on Gene Ontology annotations from using labels from January 1st, 2007 Gene Ontology. (A) Rankings labeled using January 1st, 2007 Gene Ontology annotations (solid lines) and (B) using June 22nd, 2022 Gene Ontology annotations (dashed lines).

In order to understand this performance increase on top predictions using 2022 GO labels, we analyzed the distribution of genes that had their annotations changed from 2007 to 2022. Partially due to the efforts of SPELL and MEFIT, the genes and proteins annotated to mitochondrion organization in yeast have significantly changed since 2007. One particularly interesting set is the class of 117 genes that were labeled as negatives using 2007 GO annotations (in the gold standard but not annotated to mitochondrion organization in 2007) but are labeled as positives using 2022 GO annotations (annotated to mitochondrion organization in 2022). These genes were labeled as negatives for mitochondrion organization in our training data (and in the training data of MEFIT and SPELL) but according to our current understanding of biological reality, they should be labeled as positive. Due to these mis-annotations in the Gene Ontology, these examples contaminated our negative training set. As such, these 117 “contaminated negatives” are penalized for high-ranking predictions based on the 2007 evaluation, even though their high rank is biologically accurate. As such, we analyzed the distribution of rankings for these contaminated-negative mitochondrion organization genes ranked within the top 20% of genes predicted to be involved in mitochondrion organization for each model (Fig 6). Our neural network approach ranks these reannotated genes significantly greater rank than either MEFIT or SPELL (two sample KS test, Neural network–MEFIT p-value = 0.0274; Neural network-SPELL p-value = 0.0268), suggesting that our approach is more robust to label noise in its training data, at least in this particular case.

**Fig 6.**
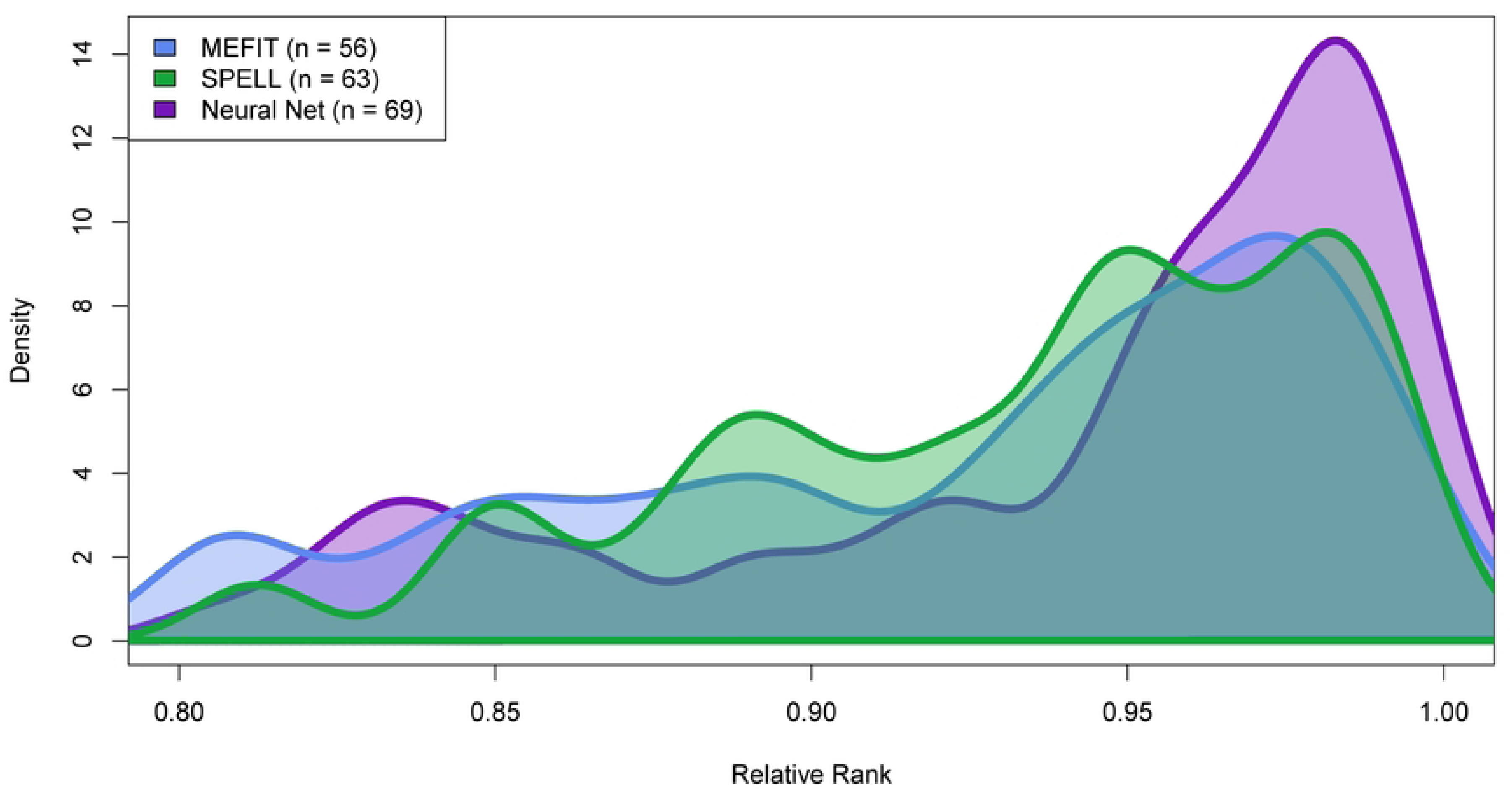
Distribution of mitochondrion organization mis-annotated negatives. Density plots of relative rank of each model of predictions of high confidence mis-annotated negative mitochondrion organization genes. A gene’s rank is calculated as its relative placement in the single gene ranking of genes involved in mitochondrion organization. For each model, n represents the number of mis-annotated negatives predicted with a relative rank between 0.8 and 1.0. Two sample right-tailed Kolmogorov-Smirov tests were performed between the distribution of the neural networks high confidence mis-annotated negatives and SPELL or MEFIT high confidence mis-annotated negative distribution. The null hypothesis was rejected for both tests (Neural networks–MEFIT p-value = 0.0274; Neural network–SPELL p-value = 0.0268).

### 3.3 Model performance using modern gene expression data and Gene Ontology labels

In the previous section, we found that our model outperformed SPELL and MEFIT when trained with the same 2007 gene expression data and Gene Ontology annotations. However, since 2007, the scientific community has accumulated significantly more gene expression data and the annotation quality of the Gene Ontology continued to improve. Therefore, we wanted to assess the performance of our approach when using modern gene expression data and Gene Ontology annotations and to explore the effects of additional input data versus improved label accuracy. Models used in this section have an architecture consisting of three layers of 80 hidden neurons each. This shared architecture was chosen for consistency of performance between the models in this section that have different sized input vectors.

We trained two models with the same 2007 Gene Ontology labels, but different gene expression input data. The first model used the 113 gene expression datasets available in 2007 and the second model used 430 gene expression datasets curated by SGD in 2022. When comparing each model’s AUC and each GO term’s single gene rankings, we found that across all GO terms, the AUC tended to be higher for the model trained with 2022 gene expression data (right-tailed paired t-test; p << 0.001) (Fig 7.A). This indicates that additional input data generally improves the accuracy of predictions.

**Fig 7.**
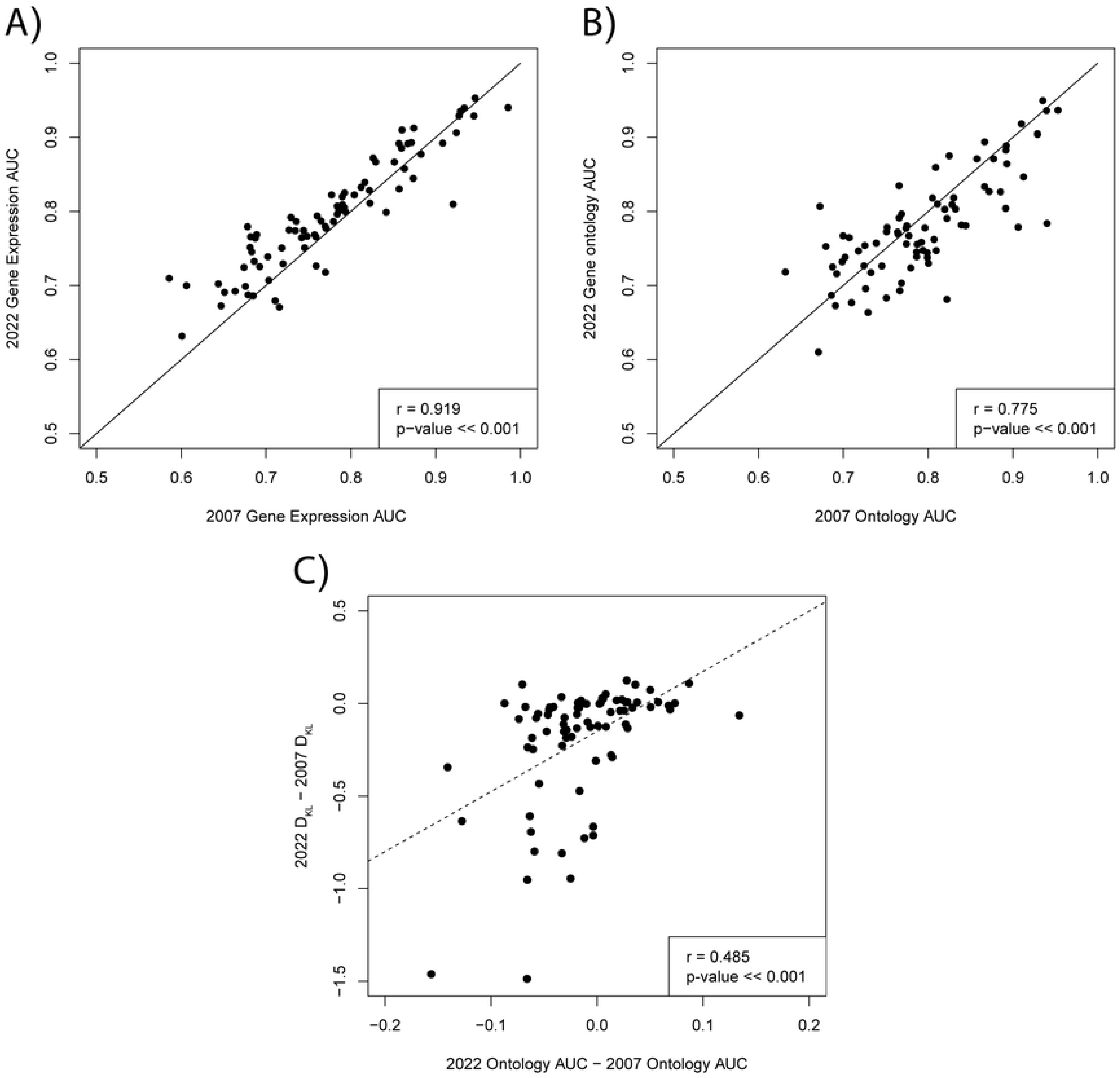
Comparison of performance with different combinations of gene expression input data and Gene Ontology training labels. (A) Scatterplot of the AUCs derived from single gene rankings for each GO term derived from two models trained with different input data and the same 2007 training labels. The x axis represents AUCs from a model trained with all gene expression datasets available in 2007 and the y axis represents AUCs from a model trained with all gene expression datasets available in 2022. The black diagonal line separates terms that perform better with 2007 vs 2022 gene expression data. (B) Scatterplot of the AUCs derived from single gene rankings for each GO term derived from two models, one trained with 2007 Gene Ontology labels (x axis) and one with 2022 Gene Ontology labels (y axis). The black diagonal line separates terms that perform better with 2007 vs 2022 Gene Ontology labels (C) For each GO term, probability distributions were created by sampling the average z-score of 200,000 genes pairs each from the set of genes annotated in 2007 (*T*_2007_) and the set of genes annotated in 2022 (*T*_2022_). Background probability distributions were created from samples of the average z-score of 200,000 gene pairs from the set of all training gene pairs in 2007 and 2022 (*B_2007_* and *B_2022_*). For each GO term, the 2007 KL divergence was measured as D_KL_(*T_2007_,B_2007_*) and the 2022 KL divergence was measured as D_KL_(*T_2022_,B_2022_*). The difference between the 2022 Ontology AUC and the 2007 Ontology AUC is plotted against the difference between the 2022 KL Divergence and the 2007 KL divergence with a regression line (dashed line).

We also trained two models using the same 2022 gene expression input data, but with different Gene Ontology training labels. The first model was trained and evaluated on labels from the 2007 Gene Ontology while the second model used labels from the 2022 Gene Ontology. We found that there was a high disparity in performance between the two models across all GO terms (Fig 7.B). Some GO terms had better AUCs when training and evaluating on 2022 labels while others performed better with the 2007 labels. Given our findings in section 3.1 that each GO term’s performance is correlated with the extent its genes are co-regulated at the level of gene expression, we decided to investigate if a similar effect could explain the disparities in performance between these models.

We hypothesized that these performance disparities could be partially explained by the change from 2007 and 2022 in how correlated the gene sets of each GO term were compared to the background distribution. For each GO term, probability distributions were created by sampling the average z-score (across 430 gene expression datasets) of 200,000 genes pairs each from the set of genes annotated in 2007 (*T*_2007_) and the set of genes annotated in 2022 (*T*_2022_). Background probability distributions were created from samples of the average z-score of 200,000 gene pairs from the set of all training gene pairs in 2007 and 2022 (*B_2007_*and *B_2022_*). For each GO term, the 2007 KL divergence was measured as D_KL_(*T_2007_,B_2007_*) and the 2022 KL divergence was measured as D_KL_(*T_2022_,B_2022_*). We found that the difference in a GO term’s AUC from 2022 labels and 2007 labels was positively correlated with the difference in the KL divergence for the 2022 and 2007 z-score probability distributions (r = 0.485; p << 0.001) (Fig 7.C). This suggests that terms that had worse performance with 2022 labels had z-score distributions that became more similar to the background distribution between 2007 and 2022, making them harder to predict.

## 4. Conclusions and future work

We created a model of automated function prediction in yeast using neural networks that can predict the functional relationship between pairs of genes in the context of 79 biological processes. Our model uses these pairwise predictions to rank individual genes by their predicted involvement for these 79 different biological processes. When training our model with the same gene expression input data and Gene Ontology labels as SPELL and MEFIT, we found that our model’s predictions of genes involved in mitochondrion organization had better ROC performance than these models. This indicates that if our model had been created at the same time as MEFIT and SPELL, it could have been used in conjunction with them to successfully direct an experimental study of mitochondrion organization [7–9]. However, our model still needs to be experimentally validated to determine if it is useful for directing experimental biology in the modern day. In future work, we plan to use petite frequency assays to test functionally uncharacterized genes our model predicts to be involved in mitochondrion organization (as did MEFIT and SPELL) [7–9].

Revisiting automated function prediction with gene expression data and Gene Ontology labels from 2007 revealed insights into how the application of neural networks for automated function prediction interacts with our accumulation of biological knowledge. We found that the accumulation of more high-throughput gene expression data between 2007 and 2022 led to a significant increase in the performance of our model across most biological processes (Fig 7.A). This suggests that despite the large corpus of gene expression data for yeast we have already amassed, putting resources into acquiring more data can still improve automated function prediction’s accuracy.

The overall conclusions for updates to Gene Ontology annotations are more complicated. We found that the difference in performance between a model trained with 2007 Gene Ontology annotations and 2022 Gene Ontology annotations varied greatly between GO terms (Fig 7.B). We found that GO terms that performed better with 2007 GO annotations had z-score distributions that were less similar to the background training distributions in 2007 compared to 2022 (Fig 7.C). These results indicate that prediction quality is highly sensitive to the annotations used to derive training labels. The quality of our training labels is restricted by the limitations of the Gene Ontology, in that GO annotations are derived from the current, incomplete knowledge curated from the scientific literature, which changes and evolves over time. Our results indicate that AFP methods must recognize these limitations when constructing training labels from GO annotations.

Our analyses of past GO annotations highlights that the way our model, and many others, define negative training examples is somewhat naive and potentially problematic. Recall that in our model, MEFIT, and SPELL, genes and pairs of genes are labeled as negative for a term if they are not annotated to that term but are annotated to another term in the GO slim.

Yeast proteins often exhibit multiple distinct functions; thus, making the assumption that a gene’s involvement in one biological process is evidence of a lack of involvement in another is reductionary. As such, there may be great value in the development of new approaches to defining negative training examples or approaches that forgo defining negative training examples entirely [33–35]. In the face of this problem, we note our model’s ability to better distinguish mis-annotated negative mitochondrion organization genes in its training set compared to SPELL and MEFIT (Fig 6). We suggest this is evidence that neural networks may have better potential to overcome the issue of problematic negatives compared to other machine learning models. However, further experiments and analyses are warranted to more fully explore this relationship.

While our model displayed better predictive accuracy than MEFIT and SPELL for mitochondrion organization, we note that our neural network model is less explainable and interpretable than these other methods. Given a predicted gene function, MEFIT and SPELL both display the input datasets that were most important for making that prediction [4,5]. Our model lacks this explainability. We did improve the interpretability of our model by finding that a GO term’s performance is correlated with the extent the co-expression of a term’s gene pairs diverge from the background training distribution (S2 Fig). That being said, our future work includes making our model more interpretable and explainable by implementing strategies like feature importance attribution [36].

We chose to build our model using linear neural networks because we wanted to compare the simplest deep learning architecture to MEFIT and SPELL. We found that linear neural networks could provide a modest increase in ROC performance over Bayesian networks and directed query searches (Fig 4). Now that we have established this baseline for deep learning models, we plan to explore other neural network architectures in our future work. In particular, we believe we can improve performance by using graph neural networks to directly learn functional relationship graphs rather than constructing them from the pairwise predictions made by our current model.

The last portion of our future work lies in testing the performance of models that integrate other modalities of biological data into its input other than just co-expression. BioPIXIE is a Bayesian network based model, similar to MEFIT, but it includes physical interaction, genomic interaction, and cellular localization data in its input. We plan to augment our model by including these data modalities in our input features [6]. Then, we want to compare this model to bioPIXIE to see if neural networks can improve performance by learning patterns between data modalities (which naive Bayesian networks are unable to do).

In summary, we have described the successful creation of a model of yeast automated function prediction using neural networks. We found that our model outperformed previous experimentally validated methodologies at predicting the involvement of yeast genes in mitochondrion organization. As such, we suspect that the predictions for other biological processes made by our model are similarly useful for directing laboratory biology experiments. Overall, we found that neural networks in their simplest form compete with or outperform other co-expression-based AFP models that use other machine learning paradigms, and changes in GO annotations and available gene expression data have a large impact on the prediction of a gene’s biological function.

## Acknowledgements

We would like to thank Dr. Bethany Strunk and Dr. Brian Teague for their help in editing the manuscript. We would also like to thank the members of Strunk Lab for their thoughtful conversations.

## Notes

### Competing Interest Statement

The authors have declared no competing interest.

